# Non-Canonical Crosslinks Confound Evolutionary Protein Structure Models

**DOI:** 10.1101/2025.03.17.643596

**Authors:** Romain Lacombe

## Abstract

Evolution-based protein structure prediction models have achieved breakthrough success in recent years. However, they struggle to generalize beyond evolutionary priors and on sequences lacking rich homologous data. Here we present a novel, out-of-domain benchmark based on sactipeptides, a rare class of ribosomally synthesized and post-translationally modified peptides (RiPPs) characterized by sulfur to *α*-carbon thioether bridges creating cross-links between cysteine residues and backbone. We evaluate recent models on predicting conformations compatible with these cross-links bridges for the 10 known sactipeptides with elucidated post-translational modifications. Crucially, the structures of 5 of them have not yet been experimentally resolved. This makes the task a challenging problem for evolution-based models, which we find exhibit limited performance (0.0% to 19.2% GDT-TS on sulfur to *α*-carbon distance). Our results point at the need for physics-informed models to sustain progress in biomolecular structure prediction.

## 1 Introduction

Evolution-based protein structure prediction models have achieved breakthrough success in recent years, leading organizers of the Critical Assessment of Structure Prediction (CASP) (Kryshtafovych et al., 2021), a longstanding bi-annual blind prediction contest, to announce the protein folding problem as “solved” for single-chain proteins (Straiton, 2023). The authors of RoseTTA Fold (Baek et al., 2021) and AlphaFold (Jumper et al., 2021) were awarded the 2024 Nobel Prize in Chemistry for their work on protein design and protein structure prediction (Nobel Prize Outreach, 2024).

Despite these major achievements, evolution-based methods, which rely on multiple sequence alignments (MSAs) or evolutionary scale pre-training, generally struggle with out-of-domain proteins for which little or no homologous data is available. They are not trained to account for the dynamics of peptide chains, which, compared to physics-based approaches, limits their ability to generalize and their performances in areas such as allosteric control, *de novo* peptide design, prediction of mutation effects, or complex interactions with other macro-molecules (Notin et al., 2023).

**Figure 1:**
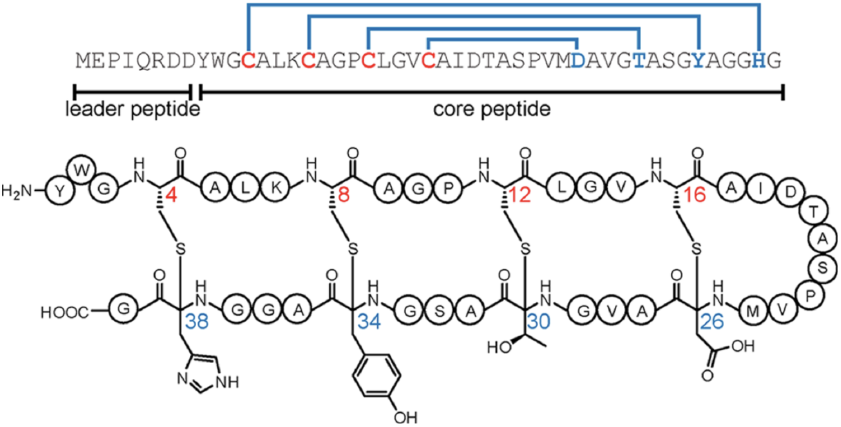
Example of sactipeptide (huazacin) and its post-translationally modified sulfur-to-alpha-carbon thioether bonds (blue nested hairpins). Figure from Hudson et al. (2019).

Evolution-based methods primarily train models on proteins with rich homology to known structures from the Protein Data Bank (PDB) (Berman et al., 2000), and benchmarks such as CASP assess them on peptides for which experimental structures have recently been resolved. Proteins with complex post-translational modifications (PTMs), especially rare ones for which little data is available, remain a challenge for structure elucidation, and therefore for protein structure prediction.

Sactipeptides are such a rare class of ribosomally synthesized and post-translationally modified peptides (RiPPs), characterized by their unique **sactionine bonds** or **‘sactibonds’**, a thioether crosslink between the sulfur in a cysteine residue and a nearby *α*-carbon atom in the backbone (Zhong et al., 2023). These post-translation modifications are crucial for the biological function of sactipeptides, yet they remain poorly represented in training data. **Their rarity makes them an ideal benchmark for testing the robustness of protein structure prediction models beyond evolutionary priors**.

In this work, we introduce a novel benchmark to evaluate the out-of-domain performance of protein structure prediction models. We define a zero-shot task: predicting the 3D cross-linking structure of sactipeptides, specifically the distance between the sulfur and *α*-carbon atoms forming sactibond bridges. We evaluate recent protein models on this task over **the only 10 known sactipeptides** with elucidated thioether cross-links, only 5 of which have an experimentally resolved structure, providing a challenging, rigorous assessment of their ability to generalize beyond their training set.

## 2 Related Works

### 2.1 Protein Structure Prediction Models

Deep learning has transformed protein structure prediction. AlphaFold2 achieved near-experimental accuracy by leveraging co-evolutionary signals through multiple sequence alignments (MSAs) (Jumper et al., 2021). RoseTTAFold introduced a three-track network that integrates sequence, MSA, and pairwise features (Baek et al., 2021). Other approaches such as ESMFold (Lin et al., 2023) and OmegaFold (Wu et al., 2022) eliminate the need for MSAs entirely by employing largescale protein language models, achieving rapid inference with competitive accuracy. Open-source models provide efficient and accessible alternatives, such as such as ColabFold (Mirdita et al., 2022), OpenFold (Bouatta et al., 2023), and Boltz-1 (Wohlwend et al., 2024).

The reliance of deep learning models on evolutionary signals still limits their applicability to proteins with poor sequence homology. Physics-based models such as Rosetta (Leaver-Fay et al., 2011) and molecular dynamics (MD) simulations (Lindorff-Larsen et al., 2011) remain essential for modeling noncanonical protein structures, intrinsically disordered regions, and cases where evolutionary priors are absent. Future progress will likely require integrating deep learning with molecular physics.

### 2.2 Benchmarking Protein Prediction

The CASP (Critical Assessment of Structure Prediction) experiment has been the gold standard for benchmarking models for three decades (Moult et al., 1995; Kryshtafovych et al., 2021). The dominance of AlphaFold 2’s in CASP14 underscored the maturity of deep learning for single-domain protein folding. However, challenges remain in modeling multi-chain complexes, alternative conformations, protein-ligand interactions, and mutation impact prediction. To address these, new benchmarks such as ProteinBench (Ye et al., 2023) and ProteinGym (Notin et al., 2023) evaluate models on broader tasks, including fitness prediction and protein design, to encourage generalization and continued progress in the field.

### 2.3 Prediction Post-Translational Modifications

Recent deep learning techniques have demonstrated impressive gains in PTM site prediction for common modifications such as phosphorylation, glycosylation, and methylation (Zhang et al., 2024; Wang et al., 2020). For RiPPs specifically, tools such as RiPPMiner (Agrawal et al., 2021) and antiSMASH (Blin et al., 2023) integrate genome mining with rule-based or machine learning pipelines to identify possible RiPP gene clusters and predict PTM topologies, bridging the gap between sequence data and experimentally validated structures.

**Table 1:** Summary of characterized sactipeptides, lengths of mature peptides, PDB entry if the structure is resolved, and post-translational cross-links structures.

| Sactipeptide | Length | PDB ID | Cross-links |
| --- | --- | --- | --- |
| Ruminococcin C1 | 44 | 6T33 | C3→N16, C5→A12, C22→K42, C26→R34 |
| Subtilisin A | 35 | 1PXQ | C4→F31, C7→T28, C13→F22 |
| Thurincin H | 31 | 2LBZ | C4→S28, C7→T25, C10→T22, C13→N19 |
| Thuricin CD $\alpha$ | 30 | 2L9X | C5→T28, C9→T25, C13→S21 |
| Thuricin CD $\beta$ | 30 | 2LA0 | C5→Y28, C9→A25, C13→T21 |
| Huazacin | 40 | — | C4→H38, C8→Y34, C12→T30, C16→D26 |
| Hyicin 4244 | 35 | — | C4→F31, C7→T28, C13→F22 |
| Skf A | 26 | — | C4→M12 |
| Streptosactin | 14 | — | C4→S7, C10→G13 |
| QmpA | 13 | — | C6→D4, C10→D8 |

## 3 Methods

### 3.1 Dataset: Sactipeptides Sequences and Cross-links

Sactipeptides—short for sulfur-to-*α*-carbon thioether-containing peptide—are a small but growing subclass of RiPPs natural products characterized by one or more intramolecular thioether linkages known as **sactionine bonds**. These unique cross-links are formed when a radical Sadenosylmethionine (rSAM) enzyme facilitates the covalent bonding of the sulfur atom of a cysteine residue to the *α*-carbon of another amino acid in the peptide backbone. The result is a tightly **cross-linked polycyclic peptide** in which the **thioether bridges structure** from residue to backbone imparts rigidity and extreme stability against heat, pH, and proteases (Flühe & Marahiel, 2013).

While the first sactipeptide—subtilosin A, an antibiotic produced by Bacillus subtilis 168—was discovered in 1985 (Babasaki et al., 1985), this class of RiPPs is still very rare, with the pace of discovery only ramping up in recent years thanks to advances in genome mining (Chen et al., 2021; Zhong et al., 2023; Wambui et al., 2022). A literature search reveals that to date, only 10 sactipeptides have a known sequence and fully elucidated cross-links structure. Of these, only 5 sactipeptides— ruminococcin C1 (Roblin et al., 2020), subtilosin A (Kawulka et al., 2004), thurincin H (Sit et al., 2011b), thuricin CD *α*, and thuricin CD *β* (Sit et al., 2011a)—have an experimentally resolved 3D structure available in the PDB. To the best of our knowledge, the remaining 5 sactipeptides— huazacin (Hudson et al., 2019), hyicin 4244 (Duarte et al., 2018), sporulation killing factor A (Cao et al., 2021), streptosactin (Chen et al., 2021), and QmpA (Ali et al., 2022)—do not have a known structure. We lists these 10 sactipeptides and their post translational cross-links in table 1.

Because half of these peptides are present in the PDB, and the other half have **identified cross-links** but ***not yet* an experimentally resolved 3D structure**, they form an ideal held-out dataset for an out-of-domain evaluation of the robustness of protein structure prediction models.

### 3.2 Cross-link Structure Prediction Task

Our goal in this work is to probe protein structure prediction models to evaluate whether they can predict a conformation consistent with post-translational cross-links observed on sactipeptides. Crucially, **the length of sulfur-to-***α***-carbon bonds is known**, measuring 1.8 Å in structures reported in the PDB. Therefore, knowing which cysteine residues form sactibonds with which target *α*-carbon atoms gives us **critical geometric information** on the 3D structure of the peptide.

We measure the predicted distances between S and C_*α*_ atoms in known sactibonds, and how closely they match this experimentally known bond length, to assess how well structures predicted by the different protein models respect the geometric bond length constraints the true structure must verify.

#### Metric: GDT-TS

The global distance test total score quantifies how many of the predicted geometries of known crosslink bonds match the experimentally observed sactibond length by less than 1 Å, 2 Å, 4 Å, or 8 Å error, with GDT-TS of 100% denoting perfect crosslink geometry prediction:

**Table 2:** Crosslinks structure prediction task results for proteins with experimentally determined structure (‘known’), and for out-of-domain sequences without a known structure (‘unknown’).

| Model | Authors | Ensembling | GDT-TS |  | RMSD |  |
| --- | --- | --- | --- | --- | --- | --- |
|  |  |  | Known | Unknown | Known | Unknown |
| AlphaFold 2 | DeepMind | Yes | 17.1 % | 7.5 % | 10.7 Å | 12.7 Å |
| AlphaFold 3 | DeepMind | Yes | 13.3 % | 11.7 % | 16.9 Å | 12.0 Å |
| Boltz-1 | MIT | No | 13.7 % | **19.2 %** | 9.7 Å | **6.9 Å** |
| ESMFold | Meta | No | 0.0 % | 10.0 % | 17.8 Å | 12.3 Å |
| OmegaFold | Tencent AI | No | 7.9 % | 9.2 % | 9.2 Å | 9.1 Å |
| RoseTTAFold 2 | Baker Lab | Yes | 16.7 % | 18.3 % | 8.1 Å | 7.9 Å |
| Average | – | – | 11.45 % | 12.65 % | 12.1 Å | 10.1 Å |

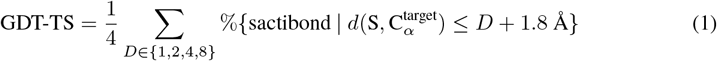

#### Metric: RMSD

This metric quantifies how far the model’s predictions are from aligning with experimentally observed sactibond geometries. The root mean square distance (RMSD) is computed between the predicted and experimentally validated sulfur-to-*α*-carbon bond distances as follows, with an RMSD of 0 Å denoting a perfect crosslink geometry prediction:

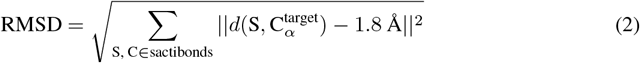

### 3.3 Experiments

We run structure prediction models for the 10 sactipeptide sequences listed in table 1 using 6 recent protein models: AlphaFold 2 (Jumper et al., 2021), Boltz-1 (Wohlwend et al., 2024), OmegaFold (Wu et al., 2022), and RoseTTA Fold 2 (Baek et al., 2023), for which we use the ColabFold MMseqs2 MSA webserver by Mirdita et al. (2022); AlphaFold 3, using the DeepMind webserver (Abramson et al., 2024); and ESM Fold, using the Meta ESM Atlas server (Lin et al., 2023). We specify random seed 42 when possible for reproducibility purposes, and use the structure with highest reported confidence for models with ensembling. We compute RMSD and GDT-TS for all models on all known sactibonds, and report the results in table 2 and figure 2.

**Figure 2:**
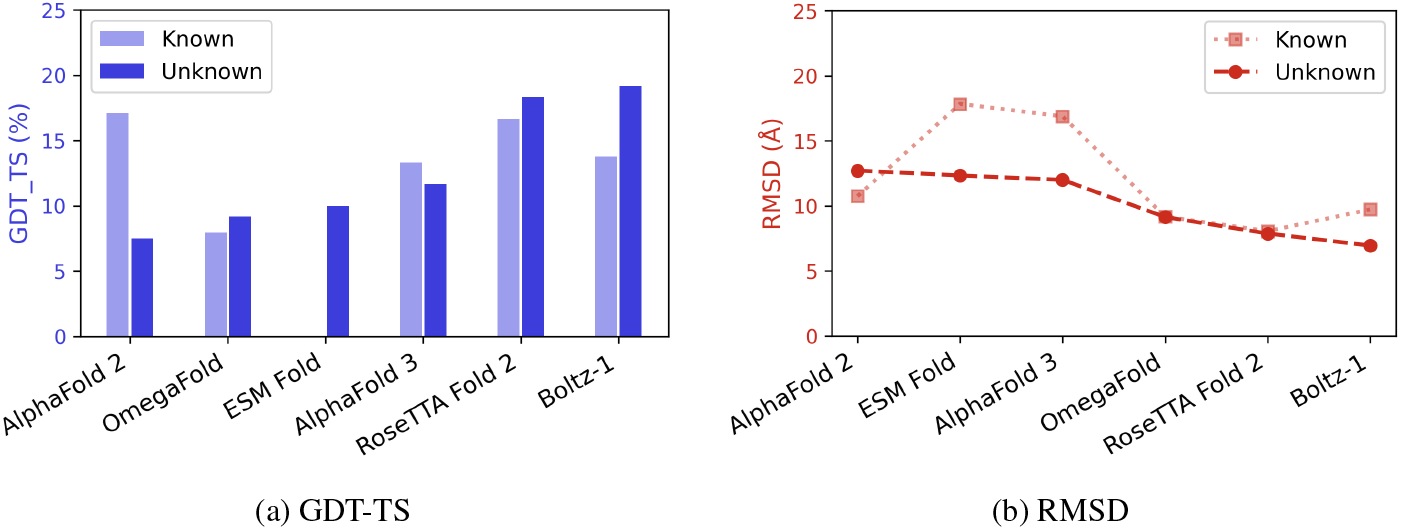
Experimental results. We report metrics for proteins with an experimentally determined 3D structure (‘known’), and out-of-domain sequences without a known structure (‘unknown’).

## 4 Results

### 4.1 Performance on Crosslink Structure Prediction Task

Our main finding is that performance on sactipeptide crosslinks structure prediction is limited for all tested models, with an average GDT-TS on known sactibonds around 11.5% and a mean RMSD of 12.1 Å (table 2). Interestingly, performance improves marginally for out-of-domain sequences with no resolved structure, with GDT-TS at 12.6% and RMSD at 10.1 Å. Boltz-1 and RoseTTAFold 2 yield the highest GDT-TS score for unknown sactipeptides, suggesting partial capture of local geometry. However, none of the models consistently approaches the ground truth 1.8 Å sulfur-to-*α*-carbon bond length, underscoring critical gaps in handling rare post-translational modifications.

### 4.2 Analysis of Predicted Structures

Visual inspection reveals that most models link sulfur atoms to one another through disulfide bonds, ignoring thioether bridge crosslinks. For example, the ruminococcin C1 structure predicted by AlphaFold 3, our highest-scoring conformation (50% GDT-TS), positions all 4 sulfur atoms in disulfide bonds. This implies heavy reliance on standard cysteine-cysteine pairing, which is interesting as the structures reported in the PDB for these sactipeptides all bear sactibonds. Models also often collapses sulfur atoms in unnatural and sterically hindered conformations, highlighting a systematic bias against thioether bonds, which we hypothesize stems from their sparsity in structural data.

**Figure 3:**
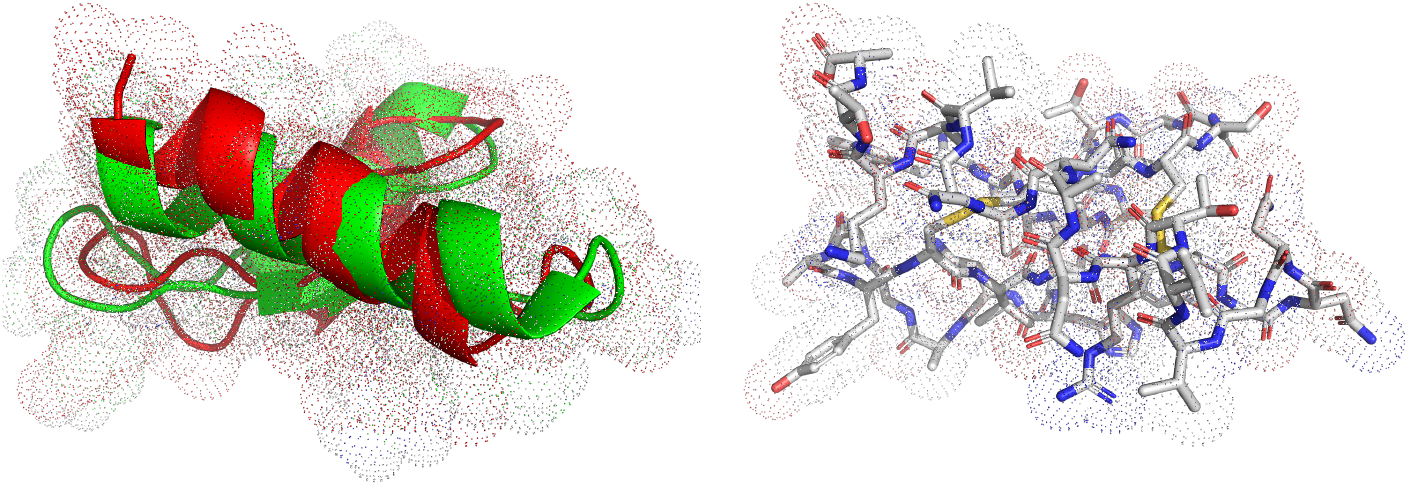
Ruminoccocin C1 structure predicted by AlphaFold 3: (a) super-imposed with the experimentally determined structure (left); (b) highlighting erroneously predicted disulfide bonds (right).

### 4.3 Limitations and Future Work

Our experiments are zero-shot and rely on pretrained models which weren’t fine-tuned on sactipeptides. Substantial improvements may therefore emerge from incorporating explicit sulfur-to*α*-carbon constraints, or from training on curated examples of these rare PTMs. We also do not explore multi-conformation sampling, which might better capture sactibond geometry. Future research could investigate fusing physics-based or data-driven potentials with evolutionary amino-acid contact maps to improve robustness and generalization.

## 5 Conclusion

We presented an out-of-domain benchmark evaluating protein structure prediction models on sulfurto-*α*-carbon thioether bonds in sactipeptides. Our findings confirm that modern evolution-based predictors, including AlphaFold and RoseTTAFold, struggle with rare post-translational polycyclic structures. Although some models perform markedly better than other, including on unknown sactipeptides, none replicate accurate crosslink bonds lengths or structure geometry. These results underscore the need for physics-informed models to better predict rare post-translational modifications, and improve predictions of the 3D structure of biomolecules beyond evolutionary priors.

## Notes

### Competing Interest Statement

The authors have declared no competing interest.

## References

Josh Abramson, Jonas Adler, Jack Dunger, Richard Evans, Tim Green, Alexander Pritzel, Olaf Ronneberger, Lindsay Willmore, Andrew J. Ballard, Joshua Bambrick, Sebastian W. Bodenstein, David A. Evans, Chia-Chun Hung, Michael O’Neill, David Reiman, Kathryn Tunyasuvunakool, Zachary Wu, Akvilė Ž emgulytė, Eirini Arvaniti, Charles Beattie, Ottavia Bertolli, Alex Bridgland, Alexey Cherepanov, Miles Congreve, Alexander I. Cowen-Rivers, Andrew Cowie, Michael Figurnov, Fabian B. Fuchs, Hannah Gladman, Rishub Jain, Yousuf A. Khan, Caroline M. R. Low, Kuba Perlin, Anna Potapenko, Pascal Savy, Sukhdeep Singh, Adrian Stecula, Ashok Thillaisundaram, Catherine Tong, Sergei Yakneen, Ellen D. Zhong, Michal Zielinski, Augustin Žídek, Victor Bapst, Pushmeet Kohli, Max Jaderberg, Demis Hassabis, and John M. Jumper. Accurate structure prediction of biomolecular interactions with AlphaFold 3. Nature, 630(8016):493–500, Jun 2024. ISSN 1476-4687. doi: 10.1038/s41586-024-07487-w. URL https://doi.org/10.1038/s41586-024-07487-w.

Priyesh Agrawal, Sana Amir, Deepak Drishtee Barua, and Debasisa Mohanty. RiPPMiner-Genome: a web resource for automated prediction of crosslinked chemical structures of RiPPs by genome mining. Journal of Molecular Biology, 433(11):166887, 2021. ISSN 0022-2836. URL 10.1016/j.jmb.2021.166887. Computation Resources for Molecular Biology.

Ataurehman Ali, Dominic Happel, Jan Habermann, Katrin Schoenfeld, Arturo Macarrón Palacios, Sebastian Bitsch, Simon Englert, Hendrik Schneider, Olga Avrutina, Sebastian Fabritz, and Harald Kolmar. Sactipeptide engineering by probing the substrate tolerance of a thioether-bond-forming sactisynthase. Angewandte Chemie International Edition, 61(45):e202210883, 2022. URL 10.1002/anie.202210883.

Katsuhiko Babasaki, Toshifumi Takao, Yasutsugu Shimonishi, and Kiyoshi Kurahashi. Subtilosin a, a new antibiotic peptide produced by bacillus subtilis 168: Isolation, structural analysis, and biogenesis1. The Journal of Biochemistry, 98(3):585–603, 09 1985. ISSN 0021-924X. URL 10.1093/oxfordjournals.jbchem.a135315.

Minkyung Baek, Frank DiMaio, Ivan Anishchenko, Justas Dauparas, Sergey Ovchinnikov, Gyu Rie Lee, Jue Wang, Qian Cong, Lisa N. Kinch, and R. Dustin et al. Schaeffer. Accurate prediction of protein structures and interactions using a three-track neural network. Science, 373(6557):871–876, 2021. URL https://10.1126/science.abj8754.

Minkyung Baek, Ivan Anishchenko, Ian R. Humphreys, Qian Cong, David Baker, and Frank DiMaio. Efficient and accurate prediction of protein structure using RoseTTAFold2. bioRxiv, 2023. URL 10.1101/2023.05.24.542179.

Helen M. Berman, John Westbrook, Zukang Feng, Gary Gilliland, T. N. Bhat, Helge Weissig, Ilya N. Shindyalov, and Philip E. Bourne. The protein data bank. Nucleic Acids Research, 28(1):235–242, 01 2000. ISSN 0305-1048. URL 10.1093/nar/28.1.235.

K. Blin, S. Shaw, H. E. Augustijn, Z. L. Reitz, F. Biermann, M. Alanjary, A. Fetter, B. R. Terlouw, W. W. Metcalf, E. J. N. Helfrich, G. P. van Wezel, M. H. Medema, and T. Weber. antiSMASH 7.0: new and improved predictions for detection, regulation, chemical structures and visualisation. Nucleic Acids Research, 51(W1):W46–W50, Jul 2023. URL 10.1093/nar/gkad344.

Nazim Bouatta, Christina Floristean, Sachin Kadyan, Ilya Baskov, Bonnie Berger, Maciej Walczak, Bryce van Allen, Gopi Diwan, Neha Kharche, and Indrani et al. Singh. OpenFold: Retraining AlphaFold2 yields new insights into its learning mechanisms and capacity for generalization. bioRxiv, 2023. URL 10.1101/2022.11.20.517210.

L. Cao, T. Do, and A. J. Link. Mechanisms of action of ribosomally synthesized and posttranslationally modified peptides (ripps). Journal of Industrial Microbiology and Biotechnology, 48(3–4): kuab005, Jun 2021. URL 10.1093/jimb/kuab005.

Yunliang Chen, Jinxiu Wang, Guoquan Li, Yunpeng Yang, and Wei Ding. Current advancements in sactipeptide natural products. Frontiers in Chemistry, 9, 2021. URL 10.3389/fchem.2021.595991.

A. F. S. Duarte, H. Ceotto-Vigoder, E. S. Barrias, T. C. B. S. Souto-Padrón, I. F. Nes, and M. D. C. F. Bastos. Hyicin 4244, the first sactibiotic described in staphylococci, exhibits an antistaphylococcal biofilm activity. International Journal of Antimicrobial Agents, 51(3):349–356, Mar 2018. URL 10.1016/j.ijantimicag.2017.06.025.

Leif Flühe and Mohamed A Marahiel. Radical s-adenosylmethionine enzyme catalyzed thioether bond formation in sactipeptide biosynthesis. Current Opinion in Chemical Biology, 17(4):605– 612, 2013. ISSN 1367-5931. URL 10.1016/j.cbpa.2013.06.031.

Graham A. Hudson, Brandon J. Burkhart, Adam J. DiCaprio, Christopher J. Schwalen, Bryce Kille, Taras V. Pogorelov, and Douglas A. Mitchell. Bioinformatic mapping of radical sadenosylmethionine-dependent ribosomally synthesized and post-translationally modified peptides identifies new cα, cβ, and cγ-linked thioether-containing peptides. Journal of the American Chemical Society, 141(20):8228–8238, 2019. URL 10.1021/jacs.9b01519.

John Jumper, Richard Evans, Alexander Pritzel, Tim Green, Michael Figurnov, Olaf Ronneberger, Kathryn Tunyasuvunakool, Russ Bates, Augustin Žídek, and Anna et al. Potapenko. Highly accurate protein structure prediction with AlphaFold. Nature, 596(7873):583–589, 2021. URL 10.1038/s41586-021-03819-2.

Karen E. Kawulka, Tara Sprules, Christopher M. Diaper, Randy M. Whittal, Ryan T. McKay, Pascal Mercier, Peter Zuber, and John C. Vederas. Structure of subtilosin a, a cyclic antimicrobial peptide from bacillus subtilis with unusual sulfur to α-carbon cross-links: Formation and reduction of αthio-α-amino acid derivatives,. Biochemistry, 43(12):3385–3395, 2004. URL 10.1021/bi0359527.

Andriy Kryshtafovych, Torsten Schwede, Maya Topf, Krzysztof Fidelis, and John Moult. Critical assessment of methods of protein structure prediction (casp)–round xiv. Proteins, 89(12):1607– 1617, 2021. URL 10.1002/prot.26237.

Andrew Leaver-Fay, Michael D. Tyka, Scott M. Lewis, Oliver F. Lange, James Thompson, Ron Jacak, Kyler W. Kaufman, P. Douglas Renfrew, Colin A. Smith, and William et al. Sheffler. ROSETTA3: an object-oriented software suite for the simulation and design of macromolecules. Methods in Enzymology, 487:545–574, 2011. URL https://10.1016/B978-0-12-381270-4.00020-8.

Zeming Lin, Halil Akin, Roshan Rao, Brian Hie, Zhongkai Zhu, Wenting Lu, Nikita Smetanin, Robert Verkuil, Ori Kabeli, and Yaniv et al. Shmueli. Evolutionary-scale prediction of atomic-level protein structure with a language model. Science, 379(6637):1123–1130, 2023. URL 10.1126/science.ade2574.

Kresten Lindorff-Larsen, Stefano Piana, Ron O. Dror, and David E. Shaw. How fast-folding proteins fold. Science, 334(6055):517–520, 2011. URL 10.1126/science.1208351.

Milot Mirdita, Konstantin Schütze, Yoshitaka Moriwaki, Lim Heo, Sergey Ovchinnikov, and Martin Steinegger. Colabfold: making protein folding accessible to all. Nature Methods, 19(6):679–682, 2022. URL 10.1038/s41592-022-01488-1.

John Moult, J.T. Pedersen, Richard Judson, and Krzysztof Fidelis. A large-scale experiment to assess protein structure prediction methods. Proteins, 23(3):ii–v, 1995. URL 10.1002/prot.340230303.

Nobel Prize Outreach. The Nobel Prize in Chemistry, 2024. URL https://www.nobelprize.org/prizes/chemistry/2024/summary/.

Pascal Notin, Aaron Kollasch, Daniel Ritter, Lood van Niekerk, Steffanie Paul, Han Spinner, Nathan Rollins, Ada Shaw, Rose Orenbuch, and Ruben et al. Weitzman. ProteinGym: Largescale benchmarks for protein fitness prediction and design. In Advances in Neural Information Processing Systems (NeurIPS) Datasets and Benchmarks Track, 2023. URL 10.1101/2023.12.07.570727.

Clarisse Roblin, Steve Chiumento, Olivier Bornet, Matthieu Nouailler, Christina S. Müller, Katy Jeannot, Christian Basset, Sylvie Kieffer-Jaquinod, Yohann Couté, Stéphane Torelli, Laurent Le Pape, Volker Schünemann, Hamza Olleik, Bruno De La Villeon, Philippe Sockeel, Eric Di Pasquale, Cendrine Nicoletti, Nicolas Vidal, Leonora Poljak, Olga Iranzo, Thierry Giardina, Michel Fons, Estelle Devillard, Patrice Polard, Marc Maresca, Josette Perrier, Mohamed Atta, Françoise Guerlesquin, Mickael Lafond, and Victor Duarte. The unusual structure of Ruminococcin C1 antimicrobial peptide confers clinical properties. Proceedings of the National Academy of Sciences, 117(32):19168–19177, 2020. doi: 10.1073/pnas.2004045117. URL https://www.pnas.org/doi/abs/10.1073/pnas.2004045117.

Clarissa S. Sit, Ryan T. McKay, Colin Hill, R. Paul Ross, and John C. Vederas. The 3d structure of thuricin cd, a two-component bacteriocin with cysteine sulfur to α-carbon cross-links. Journal of the American Chemical Society, 133(20):7680–7683, 2011a. doi: 10.1021/ja201802f. URL https://doi.org/10.1021/ja201802f. PMID: 21526839.

Clarissa S. Sit, Marco J. van Belkum, Ryan T. McKay, Randy W. Worobo, and John C. Vederas. The 3d solution structure of thurincin h, a bacteriocin with four sulfur to α-carbon crosslinks. Angewandte Chemie International Edition, 50(37):8718–8721, 2011b. URL 10.1002/anie.201102527.

Jenny Straiton. The path to solving the protein folding problem. BioTechniques, 74(5):199–201, 2023. URL 10.2144/btn-2023-0031.

Joseph Wambui, Marc J. A. Stevens, Simon Sieber, Nicole Cernela, Vincent Perreten, and Roger Stephan. Targeted genome mining reveals the psychrophilic clostridium estertheticum complex as a potential source for novel bacteriocins, including cesin a and estercticin a. Frontiers in Microbiology, 12, 2022. ISSN 1664-302X. URL 10.3389/fmicb.2021.801467.

D. Wang, D. Liu, J. Yuchi, F. He, Y. Jiang, S. Cai, J. Li, and D. Xu. Musitedeep: a deep-learning based webserver for protein post-translational modification site prediction and visualization. Nucleic Acids Research, 48(W1):W140–W146, Jul 2020. URL 10.1093/nar/gkaa275.

Jeremy Wohlwend, Gabriele Corso, Saro Passaro, Mateo Reveiz, Ken Leidal, Wojtek Swiderski, Tally Portnoi, Itamar Chinn, Jacob Silterra, Tommi Jaakkola, and Regina Barzilay. Boltz-1: Democratizing biomolecular interaction modeling. bioRxiv, 2024. URL 10.1101/2024.11.19.624167.

Ruidong Wu, Fan Ding, Rui Wang, Rui Shen, Xiwen Zhang, Shitong Luo, Chenpeng Su, Zuofan Wu, Qi Xie, Bonnie Berger, Jianzhu Ma, and Jian Peng. High-resolution de novo structure prediction from primary sequence. bioRxiv, 2022. URL 10.1101/2022.07.21.500999.

Fei Ye, Zaixiang Zheng, Dongyu Xue, Yuning Shen, Lihao Wang, Yiming Ma, Yan Wang, Xinyou Wang, Xiangxin Zhou, and Quanquan Gu. ProteinBench: A holistic evaluation of protein foundation models. arXiv preprint 2409.06744, 2023.

L. Zhang, T. Deng, S. Pan, M. Zhang, Y. Zhang, C. Yang, X. Yang, G. Tian, and J. Mi. Deepo-glcnac: a web server for prediction of protein O-GlcNAcylation sites using deep learning combined with attention mechanism. Frontiers in Cell and Developmental Biology, 12:1456728, Oct 2024. URL 10.3389/fcell.2024.1456728.

Guannan Zhong, Zong-Jie Wang, Fu Yan, Youming Zhang, and Liujie Huo. Recent advances in discovery, bioengineering, and bioactivity-evaluation of ribosomally synthesized and posttranslationally modified peptides. ACS Bio & Med Chem Au, 3(1):1–31, 2023. URL 10.1021/acsbiomedchemau.2c00062.

